# Marked differences in local bone remodeling in response to different marrow stimulation techniques in a large animal

**DOI:** 10.1101/2021.02.26.433037

**Authors:** H.M. Zlotnick, R.C. Locke, B.D. Stoeckl, J.M. Patel, S. Gupta, K.D. Browne, J. Koh, J.L. Carey, R.L. Mauck

## Abstract

Marrow stimulation, including subchondral drilling and microfracture, is the most commonly performed cartilage repair strategy, whereby the subchondral bone plate is perforated to release marrow-derived cells into a cartilage defect to initiate repair. Novel scaffolds and therapeutics are being designed to enhance and extend the positive short-term outcomes of this marrow stimulation. However, the translation of these newer treatments is hindered by bony abnormalities, including bone resorption, intralesional osteophytes, and bone cysts, that can arise after marrow stimulation. In this study, three different marrow stimulation approaches — microfracture, subchondral drilling, and needle-puncture – were evaluated in a translationally relevant large animal model, the Yucatan minipig. The objective of this study was to determine which method of marrow access (malleted awl, drilled Kirschner wire, or spring-loaded needle) best preserved the underlying subchondral bone. Fluorochrome labels were injected at the time of surgery and 2 weeks post-surgery to capture bone remodeling over the first 4 weeks. Comprehensive outcome measures included cartilage indentation testing, histological grading, microcomputed tomography, and fluorochrome imaging. Our findings indicated that needle-puncture devices best preserved the underlying subchondral bone relative to other marrow access approaches. This may relate to the degree of bony compaction occurring with marrow access, as the Kirschner wire approach, which consolidated bone most, induced the most significant bone damage with marrow stimulation. This study provides basic science evidence in support of updated marrow stimulation techniques for preclinical and clinical practice.

## Introduction

Focal cartilage injuries are common, impacting ∼ 1 million Americans annually (Mithoefer et al., 2009). Due to the poor intrinsic healing capability of articular cartilage, such lesions may progress to degenerative osteoarthritis (OA) if left untreated (Heijink et al., 2012). In the United States alone, OA impacts over 27 million people, resulting in over 100 billion dollars of healthcare costs (Jafarzadeh and Felson, 2018; Murphy and Helmick, 2012). With severe OA, joint arthroplasty may be the only option to restore mobility and improve their quality of life.

In an effort to repair cartilage defects, and ultimately prevent joint-wide OA, the most common procedure is marrow stimulation (MST) (Martín et al., 2019). MST is a relatively simple and cost-effective strategy to introduce regenerative cells into a cartilage defect. The process involves the initial debridement of the cartilage defect, followed by perforation of the subchondral bone plate using either a fluted drill bit, Kirschner wire (K-wire) (subchondral drilling) or surgical awl (microfracture) (Gao et al., 2018; Steadman et al., 2002). Pressure within the marrow compartment then promotes marrow-derived fluid and cells to populate the cartilage defect, form a clot, and serve as a template for the formation of a fibrous matrix (DiBartola et al., 2016).

While MST primarily leads to fibrous tissue formation (Knutsen et al., 2004), it remains an important first-line treatment for cartilage defect repair. MST can be performed as a single arthroscopic procedure, and has been shown to improve knee function in 70 to 95 % of patients (assessed 2-11 years post-surgery) (Gobbi et al., 2005; Knutsen et al., 2007; Steadman et al., 2003). To further improve and extend these outcomes, MST has been combined with a collagen type I/III membrane to engage and hold the marrow clot in place (Volz et al., 2017). This procedure is referred to as autologous matrix-induced chondrogenesis (AMIC), and has shown favorable results in comparison to MST alone 5 years post-surgery. In addition to AMIC, a number of laboratories have developed technologies focused on augmenting MST to direct cell differentiation and produce hyaline-like cartilage (Kim et al., 2015; Madry et al., 2020; Miller et al., 2014; Morisset et al., 2007; Patel et al., 2020; Sennett et al., 2021; Zanotto et al., 2019). Only a few of these strategies have progressed beyond the preclinical testing stage (Sharma et al., 2013).

Given the role of the subchondral bone in OA, it is likely that for augmented MST to reach its full potential, steps must be taken to address unintentional bony abnormalities that may arise from marrow access. This includes reports of bone resorption, intralesional osteophytes, and the formation of bone cysts (Cole et al., 2011; Zedde et al., 2016). These subchondral irregularities likely impede the translation of novel scaffolds and therapeutics (Sennett et al., 2021) and may also limit revision treatment options in the case that MST fails (Riff et al., 2020). Intuitively, without a stable foundation on which to form, the repair tissue may be predisposed to failure. To reduce bone damage, it may be beneficial to scale down tool sizing. Indeed, both small (1 mm) diameter awls (Orth et al., 2016) and small (1 mm) diameter K-wires were recently shown to improve MST outcomes in comparison to larger instrumentation (Eldracher et al., 2014). Others have suggested that small diameter (1 mm) holes that are deeper (9 mm depth) may preserve the underlying bone in comparison to more traditional larger (2 mm) and shallower (2 – 4 mm) holes that can fragment and compact the bone (Chen et al., 2009; Zedde et al., 2016).

While there is growing evidence to support the use of smaller diameter and deeper access for MST, the mechanism for creating these narrow, extended holes (whether it be by impaction, drilling, or another method) is not yet determined. To date, only 7 studies have directly compared the standard MST approaches – microfracture and subchondral drilling, and only 3 of these studies included multiple outcome measures (Kraeutler et al., 2020). There is thus a lack of knowledge regarding how the method of hole creation impacts the subchondral bone. Thus, the goal of this study was to evaluate the impact of 3 different methods of MST (awl-based microfracture, needle puncture, and drilling) on the underlying subchondral bone in a Yucatan minipig model of cartilage defect repair. The needle puncture device (SmartShot® Marrow Access Device) employed here is spring-powered and self-retracting, and therefore contrasts to the awl in the consistency and speed of entry and exit from the bone, and its overall penetration depth into the bone (∼4x deeper than an awl). For this study, we included one SmartShot device size matched in diameter to the microfracture awl, and another SmartShot device size matched in diameter to the K-wire, resulting in 4 experimental treatment groups for comparison. We hypothesized that the smaller SmartShot device (0.9 mm diameter) would cause the least bone resorption and compaction, as assessed 4 weeks post-MST. For this study, we also implemented bone fluorochrome labeling to track bone remodeling over the study duration. This technique was first pioneered by Drs. Berton Rahn and Stephen Perren (Rahn and Perren, 1971), and provided a unique window into the dynamics of bone healing at these MST marrow access sites.

## Materials and Methods

### Animal Study

All animal procedures were approved by the Institutional Animal Care and Use Committee (IACUC) at the University of Pennsylvania. Six skeletally mature (age 12 months at the beginning of the study) Yucatan minipigs (Sinclair Bioresources, Auxvasse, MO, USA) were used for this study. Under a single anesthetic induction, two separate surgical procedures were performed (**Fig. 1**). Both the left upper extremity and right stifle joint were sterilely draped and prepared using betadine. In the first procedure, an indwelling catheter was implanted in the brachiocephalic vein. In a second procedure, a unilateral stifle joint surgery was performed using a minimally invasive open approach (Bonadio et al., 2017). During the stifle joint surgery, a bone fluorochrome label was infused through the implanted catheter. Two weeks post-surgery, a second bone fluorochrome label was infused to track bone remodeling. Animals were euthanized 4 weeks post-surgery.

**Fig 1.**
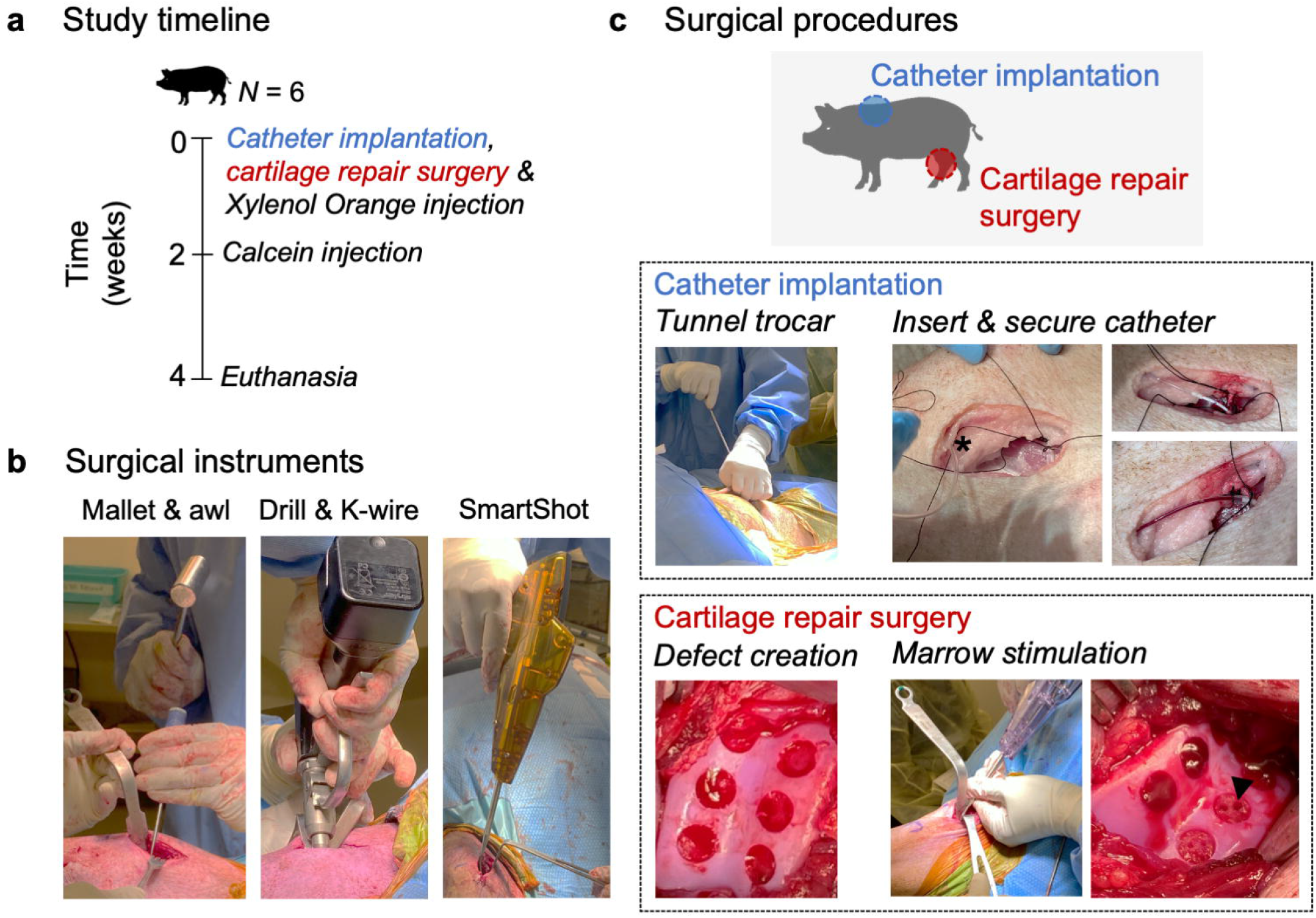
Study overview. (**a**) Study timeline. (**b**) Surgical instruments for cartilage repair procedure. (**c**) Surgical procedures. Catheter implantation: an indwelling catheter was implanted in the brachiocephalic vein for fluorochrome delivery. * denotes tunneled catheter. Cartilage repair surgery: six full-thickness chondral defects (5 mm diameter) were created in the trochlea of the stifle joint. Arrowhead: marrow stimulation hole.

#### Catheter implantation

For the catheter implantation, a 4 cm incision was made in the left neck 4.5 cm lateral to the manubrium. Dissection was carried down to the brachiocephalic vein. Medial and lateral ties (0-Silk) were loosely tied to the vein. A cannula and trocar were driven subcutaneously from the neck incision to the left scapula. The trocar was removed, and the catheter (79 cm length, Luer lock, SAI Infusion Technologies, Lake Villa, IL, USA, CPJC-12) was passed through the cannula, leaving the Luer hub exteriorized at the left scapula. An 11-blade was used to create an opening in the brachiocephalic vein for passage of the catheter. The silk ties were fastened to secure the catheter to the vein. Both the neck and scapular incisions were closed, and the catheter was flushed to assure function.

#### Stifle joint surgery

Immediately after closing the incisions from the catheter implantation, a unilateral stifle joint procedure was performed on the right hindlimb. A medial parapatellar skin incision exposed the trochlea. Six full-thickness chondral defects were created in the trochlear groove using a 5 mm biopsy punch. A curette was used to excise cartilage tissue from within the bounds of the scored defect. Care was taken to preserve the subchondral plate. In each animal, a single defect was left untreated (empty) as a control (*n* = 6). The treatment groups included: awl-based microfracture (diameter: 0.8 mm, depth: 2 mm), SmartShot (diameter: 0.9 mm, depth: 8 mm) (Marrow Access Technologies, Minnetonka, MN, USA), SmartShot (diameter: 1.2 mm, depth: 8 mm), and drilling with a Kirschner wire (K-wire) (diameter: 1.25 mm, depth: 6 mm). For each treatment, 3 marrow stimulation holes were created, except for the SmartShot (1.2 mm) where the outer housing limited the hole number to 2. The 4 treatment groups were randomized across 30 defects (*n* = 7-8/treatment group).

#### Fluorochrome injections and catheter maintenance

Xylenol Orange (90 mg/kg, MilliporeSigma, Burlington, MA, USA, 398197) was injected slowly over 1 hour via the indwelling catheter during the stifle joint procedure (Funk et al., 2009). Xylenol Orange was dissolved in 100 mL saline (Medline, Wilmer, TX, USA, EMZ111240) and sterile filtered. 2 weeks post-surgery, the animals were sedated, anesthetized, and intubated. Under anesthesia, Calcein (15 mg/kg, MilliporeSigma, C0875) was injected slowly over 1 hour via the indwelling catheter (Funk et al., 2009). Calcein was dissolved in 100 mL of a 1.4 % (wt/vol) sodium bicarbonate (MilliporeSigma, S5761) solution and sterile filtered. Blood draws were performed via the catheter before each fluorochrome injection, 1 minute post-injection, 1 day post-injection, and 1 week post-injection. Blood samples were analyzed for ionized calcium (Antech Diagnostics). The catheters were flushed twice daily with saline, and Heparin locked (100 U/mL, Medsupply Partners, Atlanta, GA, USA, BD-306423) to maintain patency. All animals wore Thundershirts for dogs (Thunderworks, Durham, NC, USA, BSSXXL-T01) to protect the exteriorized catheter hub and prevent pull-out.

#### Euthanasia and cadaveric use of non-operative stifle joint

All animals were euthanized 4 weeks post-surgery. The operative limbs were prepared for cartilage indentation testing. Four of the non-operative contralateral stifle joints (left hindlimbs) were used to assess initial (time zero) local bone compaction surrounding the marrow stimulation holes. For this assessment, mock surgeries were performed on the cadaveric joints, mimicking the operative limbs. Briefly, chondral defects (*n* = 6 defects/knee) were created using a 5 mm biopsy punch and curette. Each limb had 1 non-treated (empty) defect control (*n* = 4 empty defects), and the surgical treatments (microfracture (0.8 mm), SmartShot (0.9 mm), SmartShot (1.2 mm), K-wire (1.25 mm)) were randomized to the remaining defects (*n* = 5 defects/treatment).

### Cartilage indentation testing

Following euthanasia, a hand saw was used to produce 1 cm^3^ osteochondral units containing the chondral defect from the operative limbs (Sennett et al., 2021). Control cartilage was collected from the distal end of the trochlea. Each sample was potted in polymethyl methacrylate (PMMA) (Ortho-Jet, Lang Dental, Wheeling, IL, USA) to secure the bone. Samples were covered in phosphate buffered saline (PBS) with protease inhibitors (Roche Complete, MilliporeSigma) and tested within 5 h of collection. Each sample was place in a custom rig equipped with an XY positioning stage and a goniometer to ensure that the cartilage surface was indented perpendicular to the 2 mm diameter spherical indenter (Meloni et al., 2017). A load of 0.1 N was applied at an initial rate of 0.1 mm/s and held over 900 s as the tissue underwent creep displacement. The compressive modulus (E_y-_) was computed by fitting the creep data to a Hertzian Biphasic analytical model (Moore et al., 2016). Cartilage thickness was experimentally calculated from samples excised during surgery, and input into the analytical model.

### µCT

After indentation testing, samples were removed from the PMMA, submerged in PBS with protease inhibitors, and imaged via µCT (Scanco µCT50, Scanco Medical, Southeastern, PA, USA). Scans were conducted utilizing the following parameters: 3000 projections, 900 ms x 5 exposures/projection, voltage: 45 kVp, current: 133 µA, isotropic voxel size: 4.4 µm. The volume of bone resorption was calculated using the Scanco evaluation software by manually contouring the void space underneath the neocartilage. The marrow stimulation holes were excluded from these measurements.

Parallel µCT scans were performed on the osteochondral units from the cadaveric time zero surgeries using the contralateral hindlimbs. The same µCT imaging parameters were used for these samples. For each treatment condition, the local bone volume/total volume surrounding individual marrow stimulation holes (*n* = 6 holes analyzed/treatment) was calculated. Using the Scanco evaluation software, an outer circle was drawn with a diameter 0.65 mm greater than the original hole diameter. This value was selected to ensure that the volume measured was unique to that hole, and not overlapping with another marrow stimulation hole within the same defect. Subsequently, the perimeter of the marrow stimulation hole was contoured within this outer circle, creating a hollow cylinder. The bone volume/total volume was computed for this hollow cylinder, excluding the inner void space.

### Histology

#### Mineralized cryohistology

Samples were fixed immediately after µCT scanning for 24 h in 10 % neutral buffered formalin. After fixing, the samples were infiltrated with a 10 % sucrose (Fisher Scientific, S25590) and 2 % polyvinylpyrrolidone (MilliporeSigma, P5288) solution for 48 h. Samples were then embedded in optimal cutting temperature compound (OCT), and sectioned (18 µm/section) undecalcified with cryofilm until reaching the midplane of the defect (Dyment et al., 2016). Tape-stabilized, frozen sections were subjected to two rounds of imaging on a Zeiss Axio Scan.Z1 digital slide scanner, including imaging of (i) fluorochrome labels, and dark field, and (ii) tartrate-resistant phosphatase (TRAP) staining with TO-PRO-3 Iodide (ThermoFisher Scientific, T3605) counterstain. For TRAP staining, sections were incubated in TRAP buffer (0.92% sodium acetate anhydrous, 1.14% _L-(+)_-tartaric acid, 1% glacial acetic acid – pH 4.1-4.3) for 1 h, and then incubated with ELF97 substrate (ThermoFisher Scientific, E6588) in buffer for 1 h under ultraviolet light.

The fluorochrome labels (% area) were quantified in the region surrounding the marrow stimulation holes. Images were split into red and green channels, representing the xylenol orange, and calcein signals, respectively. The images were then binarized, and rotated in FIJI to orient the marrow stimulation holes perpendicular to the surface (Schindelin et al., 2012). A region of interest (3 mm by 1 mm) was defined immediately adjacent to the marrow stimulation hole. The TRAP staining was quantified by measuring the number of positive trap pixels within a 7 mm by 7 mm area, normalized by the bone area within this region.

#### Demineralized paraffin-wax histology

After cryosectioning, one half of each defect, the samples were removed from OCT and decalcified (Formical 2000, StatLab Medical Products, Columbia, MD, USA) for 2 days. These decalcified samples were then processed for paraffin-wax histology. Sections from the midplane of each defect were cut (7 µm), and stained with haematoxylin and eosin and safranin O/fast green (SafO/FG).

#### Histological scoring

Safranin-O/Fast Green (SafO/FG) stained sections were assessed using the modified ICRS II scoring system (Mainil-Varlet et al., 2010). The following parameters were scored on a continuous scale from 0 (poor defect repair) to 100 (normal articular cartilage): defect fill, integration to surrounding cartilage, matrix staining, surface architecture, basal integration, subchondral bone abnormality, vascularization, surface/superficial assessment, mid/deep zone assessment, overall assessment. Three blinded reviewers with expertise in cartilage histomorphology scored each sample. Scores for each defect were averaged and plotted by treatment group.

### Data analysis

All quantitative data were analyzed with GraphPad Prism (version 8.4.3 for MacOS, GraphPad Software, San Diego, CA, USA). For mechanical testing, histological scoring, blood analyses, fluorochrome labeling, bone resorption, and TRAP data, a one-way analysis of variance (ANOVA) was performed with post-hoc Tukey’s corrections for multiple comparisons. For the bone compaction analysis, a Kruskal Wallis test was performed a with post-hoc Dunn’s test. All plots were made in Prism.

## Results

### Surgical outcomes and blood analyses

All animal procedures were performed without complications (**Fig. 1**), and animals were weight bearing within 2 h post-surgery. Animal skin coloration changed dramatically immediately after fluorochrome injection (**Fig. 2a**), evidence of the systemic distribution of label. The xylenol orange and calcein injections caused the animals to transiently turn purple and yellow, respectively. Animal skin coloration returned to normal within a few hours post-injection. Blood samples were taken both pre- and post-injection (1 minute, 1 day, 1 week), and analyzed for ionized calcium levels. There was a slight decrease in ionized calcium after the xylenol orange label (**Fig. 2b**), but this returned to baseline within 1-week post-injection. After the calcein injection, there were no changes in ionized calcium (**Fig. 2c**).

**Fig 2.**
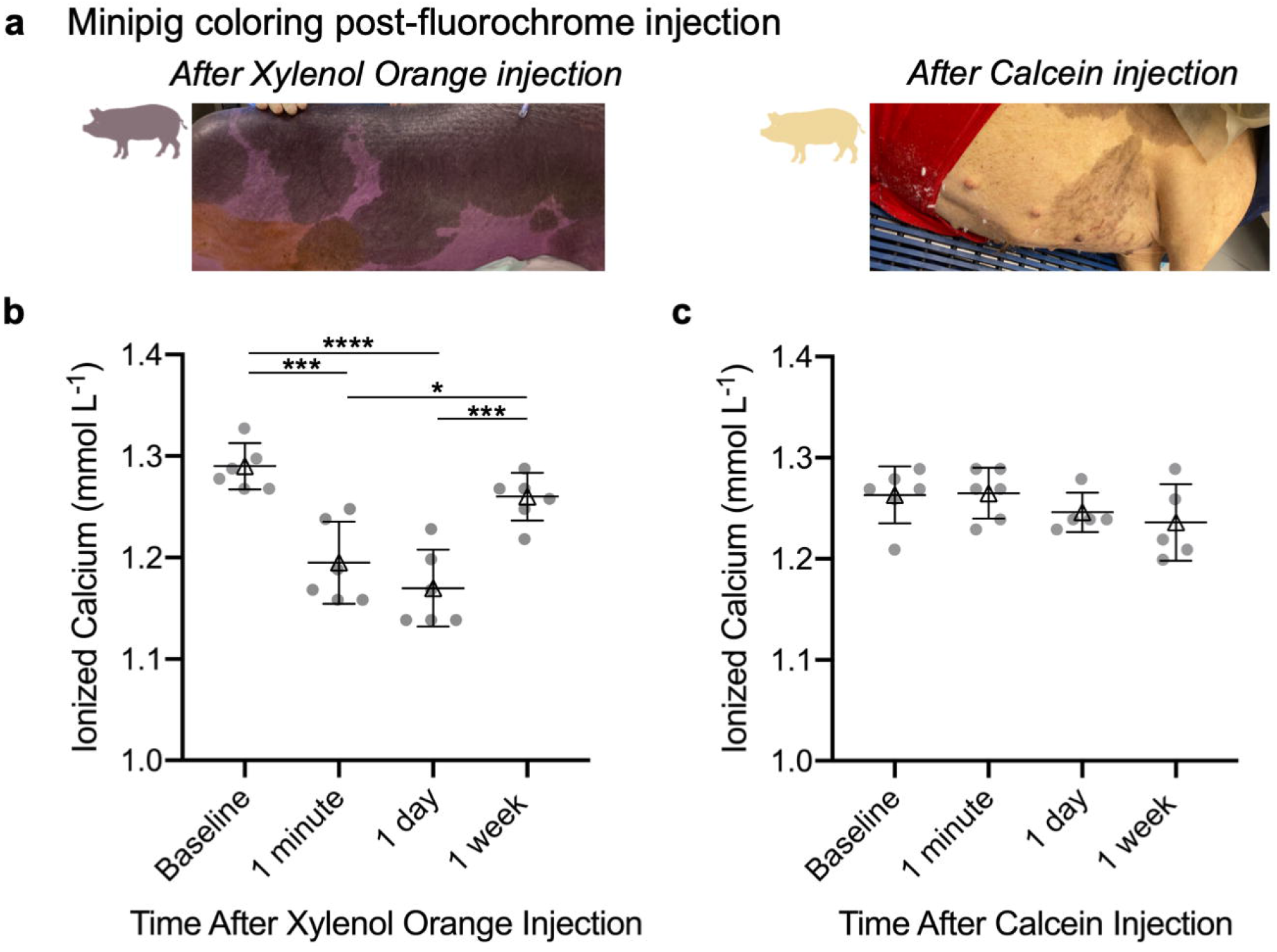
Blood analyses before and after injecting fluorochrome labels. (**a**) Minipig coloring post-fluorochrome injection. Immediately after the xylenol orange injection, the animals turned purple. The animals turned slightly yellow after the Calcein injection. Animal coloring returned to normal within 3 h post-injection. (**b, c**) Ionized calcium levels in the blood after xylenol orange (b) and calcein (c) injection through 2 weeks post-surgery. For both plots, individual animal data is represented by the circles. The triangles and error bars represent the mean ± standard deviation. \**p* < 0.05, \*\*\**p* < 0.001, \*\*\*\**p* < 0.0001. N = 6 animals.

### Mechanical testing and Histology of Regenerating Cartilage

Indentation testing was performed to assess the mechanical properties of the repair tissue (**Fig. 3a**). Native cartilage controls were taken from the distal portion of the operative femoral trochlea. As expected, all treatment groups had a significantly lower compressive modulus in comparison to the native cartilage controls (**Fig. 3b**). There were no differences observed between marrow stimulation groups at this early time point (**Fig. 3c**). Similarly, histological assessment showed similar cartilage repair between treatment groups 4 weeks post-surgery. Faint outlines of the original marrow stimulation holes were present in both SmartShot conditions, whereas more prominent void spaces were visible in the K-Wire group (**Fig. 4a**). Blinded ICRS II scoring revealed improved basal integration in the 0.9 mm SmartShot group compared to the microfracture group (**Fig. 4b**). There were no significant differences between treatment groups in the nine other parameters scored: defect fill, integration to surrounding cartilage, matrix staining, surface architecture, subchondral bone abnormality, vascularization, surface/superficial assessment, mid/deep zone assessment, and overall assessment. As evident from the representative SafO/FG images, matrix staining was minimal across all groups.

**Fig 3.**
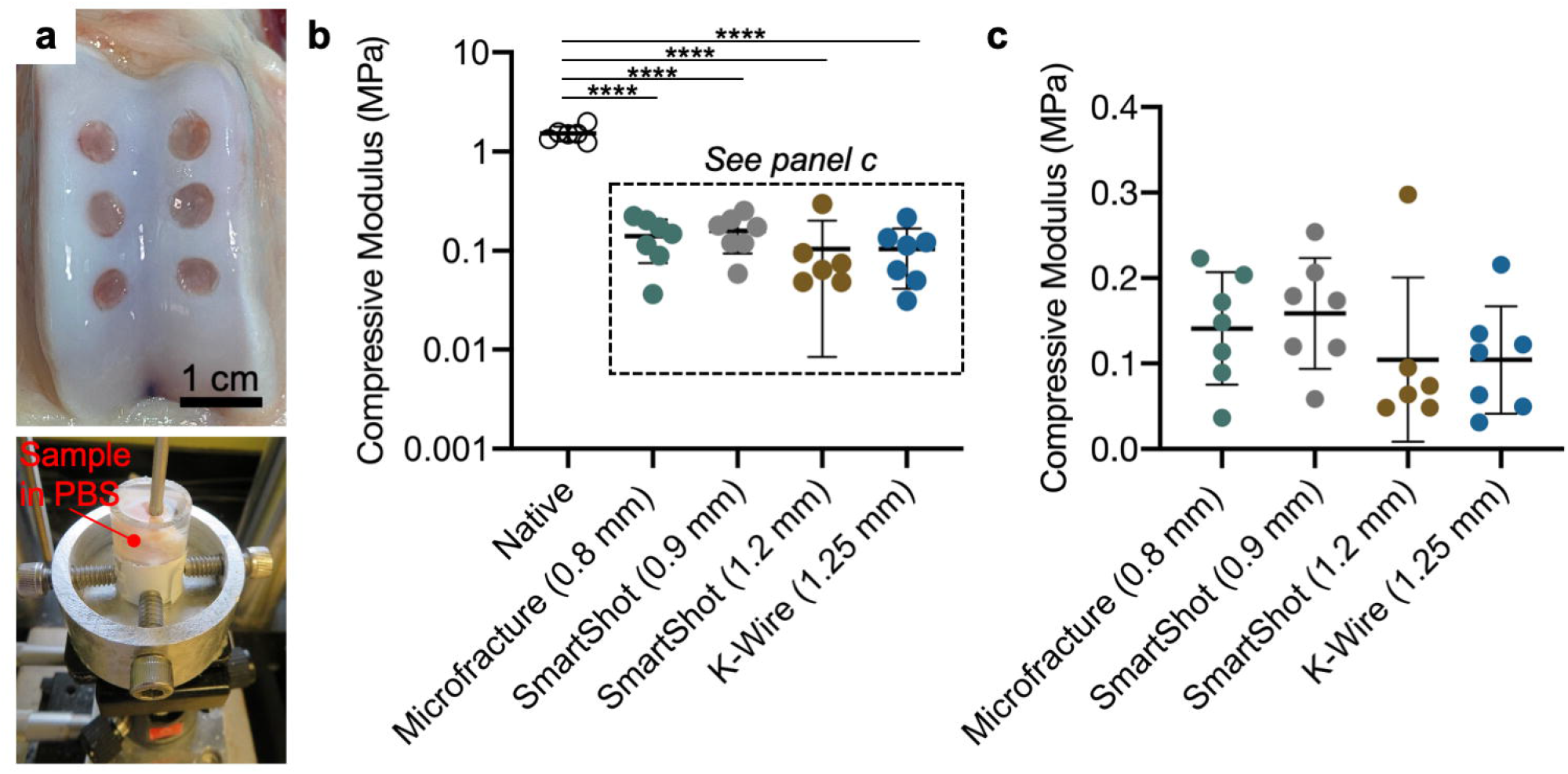
Macroscopic cartilage repair and cartilage mechanical properties. (**a**) Macroscopic image of joint 4 weeks post-surgery, and indentation testing setup. (**b**) Compressive modulus (displayed on log_10_ scale) including the native cartilage controls. \*\*\*\**p* < 0.0001. (**c**) Compressive modulus (linear scale, from b). All plots show mean standard ± deviation. n = 7 defects/treatment. N = 6 animals.

**Fig 4.**
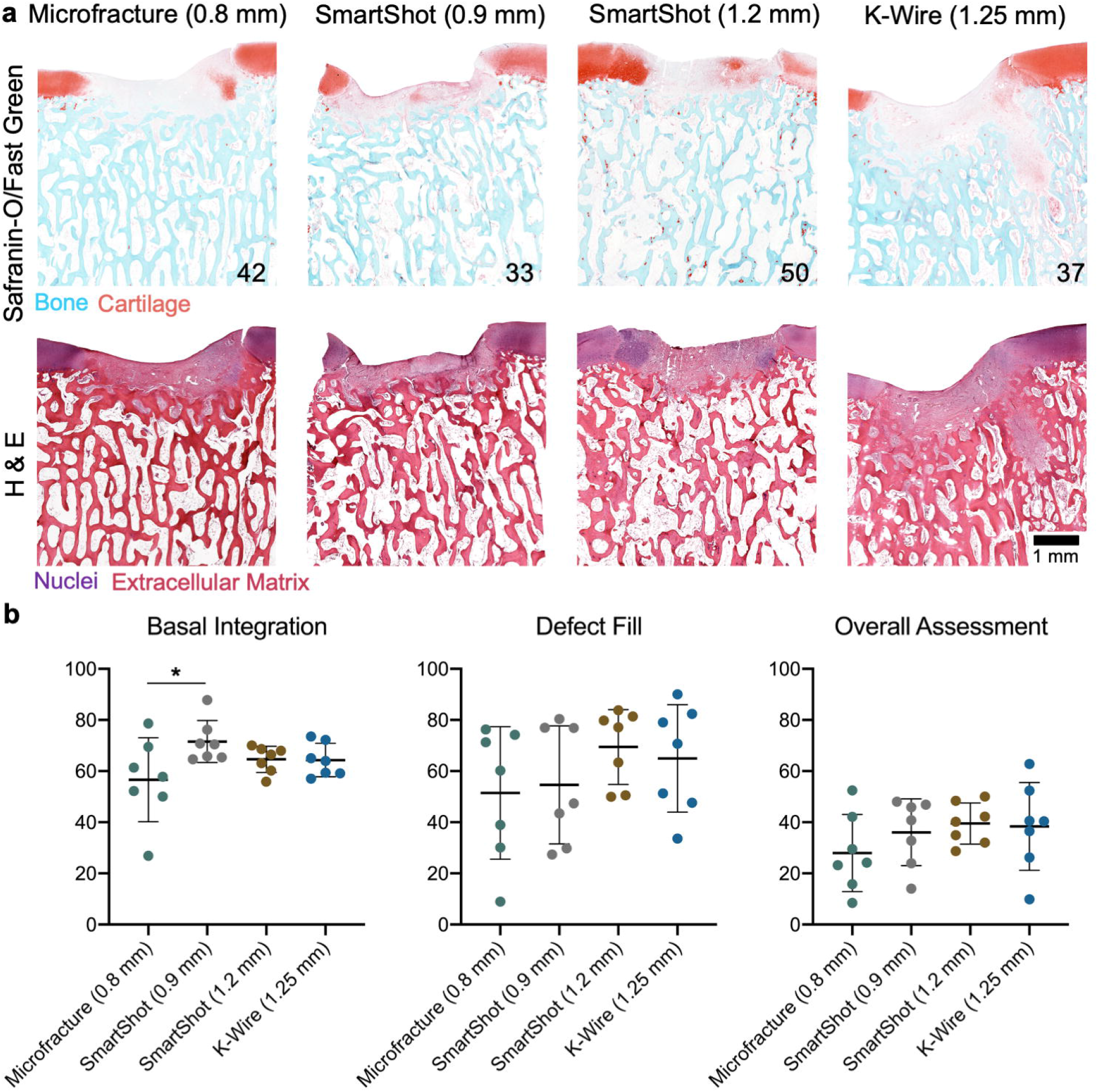
Cartilage repair quality 4 weeks after MST. (**a**) Histological staining (Safranin-O/Fast Green and Hematoxylin/Eosin) of representative defects. Numbers represent the overall ICRS II histological score for that specimen. Limited proteoglycan deposition was observed in the defects at this early time point. (**b**) Blinded ICRS II histological scoring (0: worst, 100: best). \**p* < 0.05. Bars indicate mean ± standard deviation. n = 7 defects/treatment. N = 6 animals.

### Fluorochrome imaging

Fluorochrome label incorporation at the time of surgery (xylenol orange), and 2 weeks post-surgery (calcein) showed marked differences in the bony remodeling response from each marrow stimulation technique (**Fig. 5a**). For this analysis, a region of interest was set adjacent to the marrow stimulation hole (**Fig. 5b**). The percent area of each mineral label was quantified within this region of interest. The defects treated with the SmartShot devices had the highest xylenol orange incorporation, indicative of early bone formation, while a delayed response was seen in the K-Wire group (**Fig. 5c**). At the later time point (2 weeks), the K-Wire group had significantly higher calcein incorporation in comparison to all other groups (**Fig. 5d**). The defects treated with microfracture awl had relatively even percentages of each label, signifying a steady, but relatively quiet, bony response over the 4 weeks testing period.

**Fig 5.**
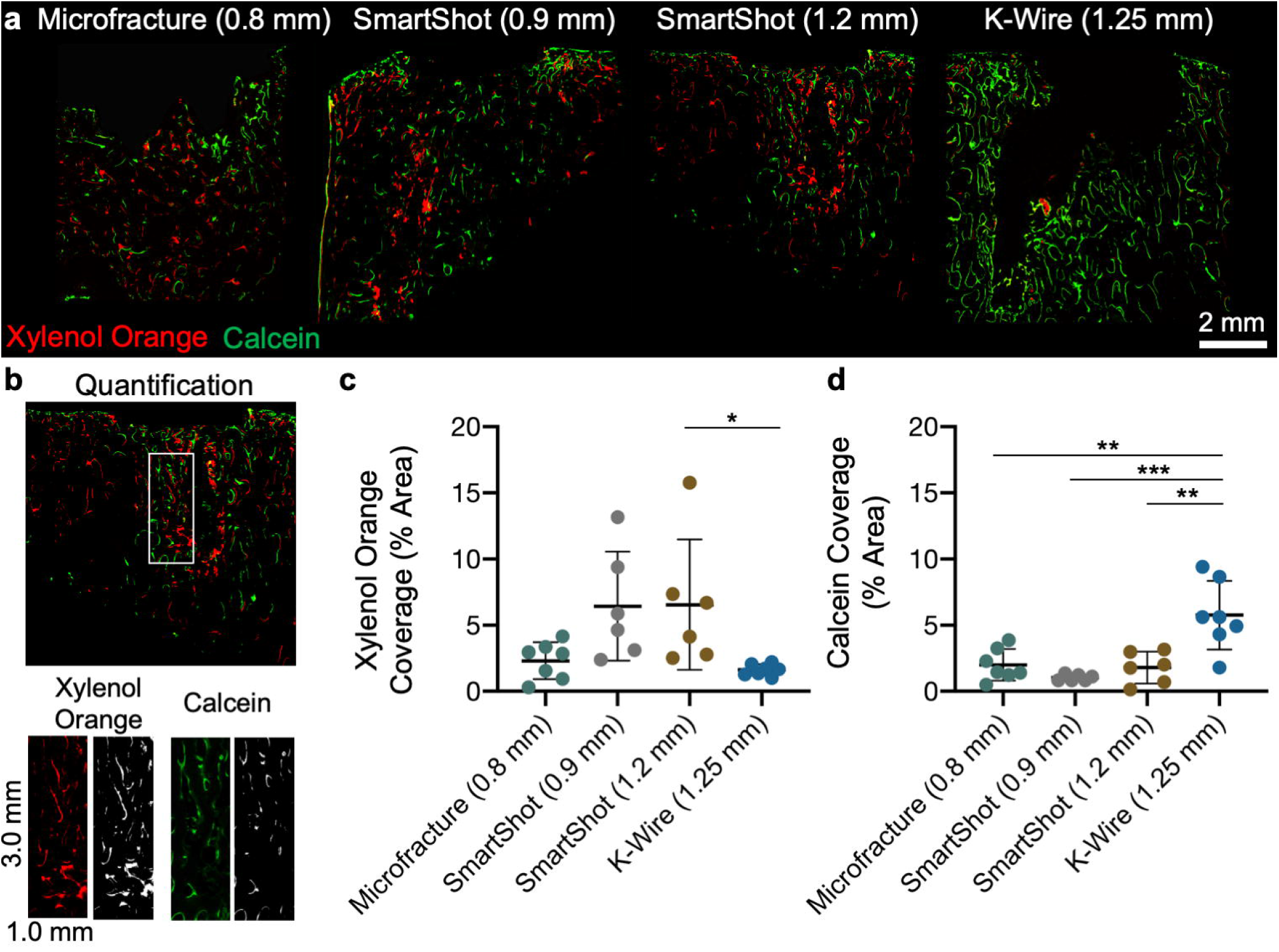
Bone remodeling over 4 weeks after MST. (**a**) Representative fluorochrome images at the midplane of the defects. Red: Xylenol Orange, injected at the time of surgery. Green: Calcein, injected 2 weeks post-surgery. (**b**) Quantification of fluorochrome images. A 3.0 mm by 1.0 mm rectangular region nearest to the marrow stimulation site was separated into the respective red and green channels, and quantified. (**c, d**) Percent area of Xylenol Orange and Calcein, respectively. \**p* < 0.05, \**\*p* < 0.01, \*\*\**p* < 0.001. Bars indicate mean ± standard deviation. n = 6-7 holes/treatment. N = 6 animals.

### Global bony response

µCT was used to assess bone resorption in the subchondral bone underlying the cartilage defects 4 weeks post-surgery (**Fig. 4a**). Similar to the histology findings, the marrow stimulation holes were most apparent in the K-Wire group. For each treatment group, the marrow stimulation holes were excluded from the bone resorption measurements. 3D quantification of bone resorption indicated that the K-Wire group had significantly more bone resorption compared to the 1.2 mm SmartShot. Cryosections stained for TRAP, which is expressed by osteoclasts, revealed that there was higher osteoclastic activity in the subchondral region of defects treated with the K-Wire in comparison to both the 1.2 mm SmartShot and native bone.

### Bone compaction

Immediately after euthanasia, the contralateral hindlimbs were used to investigate the initial bone compaction caused by each marrow stimulation device (**Fig. 7a**). For this time zero assessment, we created chondral defects, randomized the treatment groups across defect locations, and performed µCT imaging on the osteochondral units after marrow access. Here, higher bone volume/total volume (BV/TV) measurements represented increased bone compaction. The K-Wire group showed a significantly higher BV/TV in the region immediately adjacent to the marrow access holes in comparison to both the 0.9 mm SmartShot device and native bone. The range of BV/TV values measured from each SmartShot device overlapped with the native bone BV/TV range, unlike the microfracture and K-Wire groups.

**Fig 6.**
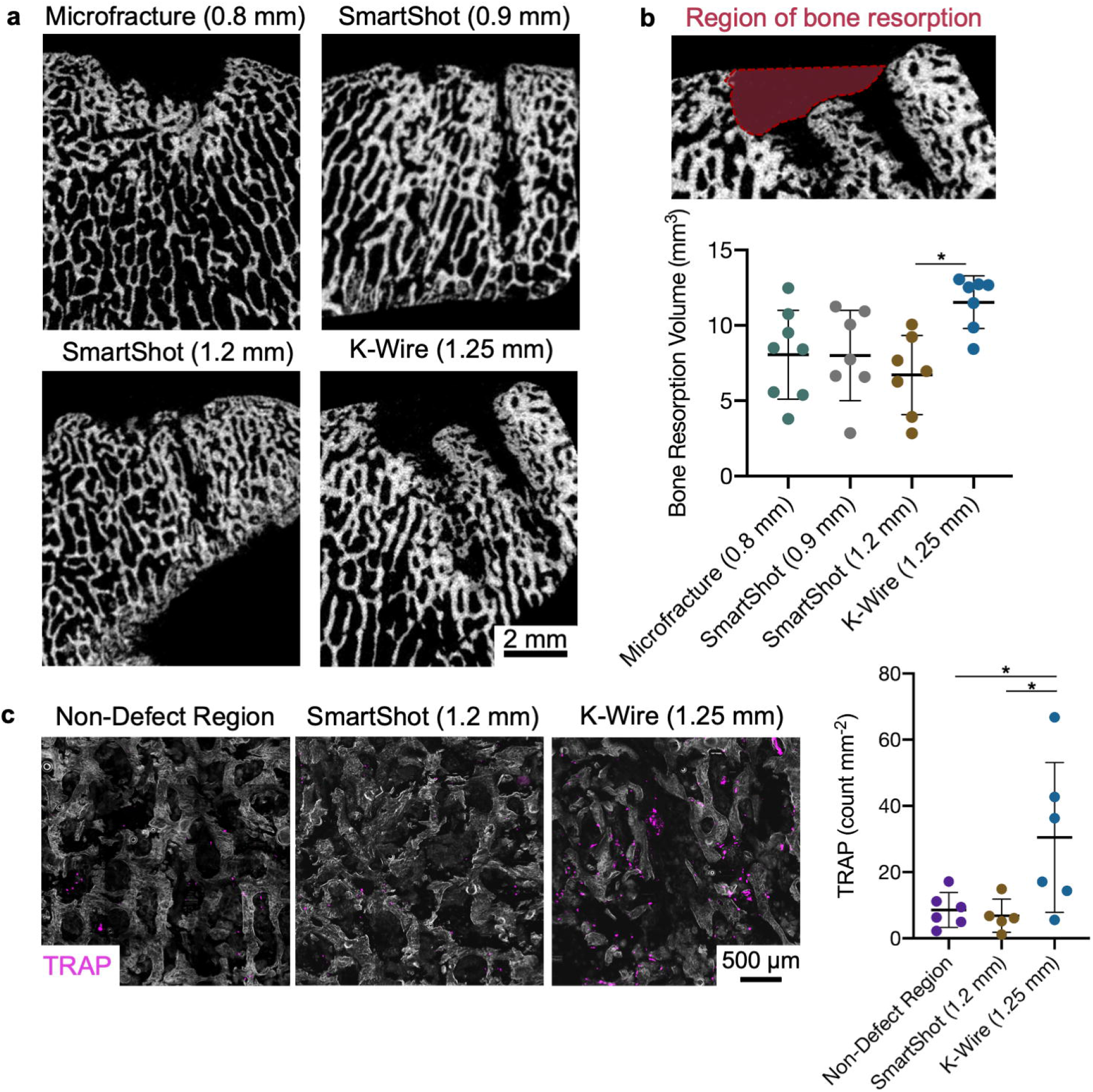
Global bony response underneath cartilage defects following MST. (**a**) Representative µCT images at the midplane of the defects. (**b**) 3D quantification of bone resorption. \**p* < 0.05. n = 7-8 defects/treatment. N = 6 animals. (**c**) Tartrate-resistant phosphatase (TRAP) staining and quantification. TRAP positive pixels were normalized to the bone area in a 7 mm by 7 mm image. \**p* < 0.05. Bars indicate mean ± standard deviation. n = 5-6 defects/treatment. N = 6 animals.

**Fig 7.**
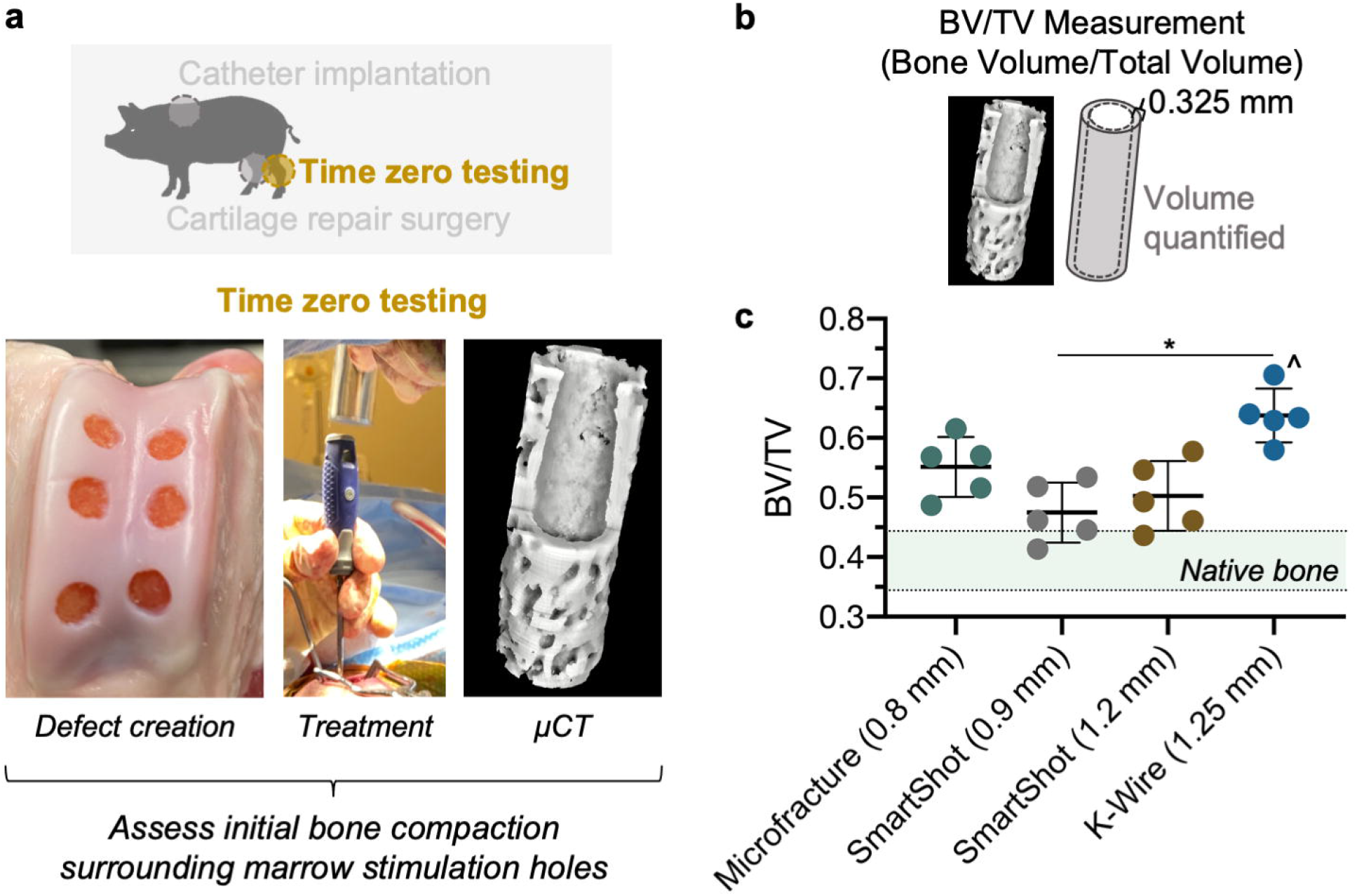
Bone compaction surrounding fresh (time zero) marrow stimulation hole. (**a**) Schematic of time zero testing. (**b**) Bone volume/total volume (BV/TV) in region surrounding marrow stimulation hole. (**c**) Native bone was measured underneath a cartilage defect without any marrow stimulation holes, and the range of values are displayed by the shaded region. \**p* < 0.05. ^ *p* < 0.001 in comparison to native bone. Bars indicate mean ± standard deviation. n = 5 holes/treatment. N = 4 animals.

## Discussion

This study directly compared bony changes that occur post-MST using three different methods of marrow access hole creation—awl-based microfracture, needle puncture (SmartShot), and subchondral drilling (using a K-wire). Although not expected to show differences, this study also characterized the cartilage repair quality from each of these treatments at the 4-week terminal timepoint. Comprehensive analyses were performed to assess bone quality post-MST including, histology, µCT, and fluorochrome labeling. Taken together, these results highlight potential disadvantages in the use of K-wires for subchondral drilling. The 1.25 mm K-wire caused the most bone resorption, delayed bony healing, and compacted the bone around the perimeter of the MST holes in comparison to other marrow access methods. It may be that a fluted drill bit (Gao et al., 2018), designed to excise bone chips as the drill advances, would better preserve the subchondral bone, in comparison to the smooth K-wire, however that was not assessed in this study. A smaller diameter K-wire may have also caused less damage to the bone. Regardless, the 1.25 mm K-wires used in this study are within the range commonly used in clinical practice. As an alternative approach, the SmartShot device best preserved the subchondral bone. Interestingly, we found no statistical differences between the different sized SmartShot devices, suggesting that the method of hole creation may play a larger role than hole size in terms of the post-MST bony response.

Previous studies have highlighted the variability in MST, from the technique itself (Kroell et al., 2016; Theodoropoulos et al., 2012) to patient outcomes (Knutsen et al., 2007; Mithoefer et al., 2009). There is considerable intra- and inter-surgeon variability with MST (Kroell et al., 2016), which likely contributes to the variable cartilage fill grade (18 to 95%)(Mithoefer et al., 2009). While the main focus of this study was not on improving the repeatability of MST, our results do support the use of the SmartShot (or similar device), which could offer greater control and uniformity of hole depth across surgeons. Further, the use of this device in both the preclinical and clinical procedures may enhance the repeatability of outcomes between research groups and surgical teams.

One of the primary advances of this study was the implementation of fluorochrome labeling in a large animal. This is the first known study that used fluorochrome imaging to monitor the dynamic bone remodeling that occurs after marrow stimulation techniques. This longitudinal outcome measure reduces the number of animals required to visualize bony changes over time. While others have performed fluorochrome labeling in minipigs (Funk et al., 2009), this technique is relatively uncommon in large animals. The scarcity of large animal studies utilizing fluorochrome labeling may be due to the fact that while such mineral labels can be injected subcutaneously into small animals, large animals require intravenous administration (van Gaalen et al., 2010). Additionally, many of the fluorochrome labels used in rodents are not compatible with larger animals. Specifically, alizarine complexone can lead to sudden death in minipigs (Ruehe et al., 2008). To ensure animal safety over the duration of this study, we took periodic blood samples from the intravenous catheter, and analyzed the blood samples for ionized calcium. While there were statistically significant differences in ionized calcium levels after injecting xylenol orange, these transient changes are likely not physiologically significant (that is, ionized calcium remained in the physiologic range). Overall, the positive response of our animals to the described fluorochrome labeling protocol should encourage other groups to implement this potentially impactful outcome measure in studies of MST and osteochondral regeneration.

All large animal studies are limited by sample size; nonetheless, we demonstrated an efficient utilization of animals, and were able to identify significant differences between treatment groups within our cohort, demonstrating the utility of study design to reduce animal number. Both the operative and non-operative hindlimbs were used to establish two time points (time zero, and 4 weeks) for separate outcome measures. Additionally, both the cartilage repair quality, and changes to the subchondral bone were assessed. While we did not expect to detect differences in cartilage repair at the 4-week time point (given how short the time period was), it was important to perform such assays to provide a baseline and motivation for future long-term studies evaluating next-generation cell and material adjuvants to MST. Future work will also determine whether bone quality relates to repair tissue strength and durability.

## Conclusions

This study built on previous literature suggesting that small diameter and deeper MST holes may reduce subchondral bone abnormalities and ultimately improve cartilage repair. While a few studies have compared tool sizing for MST, none have jointly investigated tool sizing and the method of hole creation. This study provides strong evidence in favor of a repeatable needle-puncture device (SmartShot or similar device) to best preserve the subchondral bone and minimize aberrant remodeling after MST. This work also provides a cautionary tale regarding the use of K-wires, as this treatment group consistently performed poorly with respect to subchondral bone remodeling and healing. Overall, the results of this study should inform the design of future preclinical studies, and has the potential to change current surgical practice.

## Acknowledgements

This study was supported by the NIH (R01 AR077362, T32 AR007132, P30 AR069619, F31 AR077395). Additional funding was provided by the Department of Veterans’ Affairs, and the NSF (CMMI: 15-48571). The authors are grateful for the University of Pennsylvania’s University Laboratory Animal Resources (ULAR) veterinary staff. Additionally, the authors would like to acknowledge Drs. Sarah Gullbrand, Sherry Liu, Thomas Schaer, Julia Funk, and Chelsea Wallace for their experimental guidance and advice. Lastly, the authors would like to thank Marrow Access Technologies for donating the SmartShot devices, and Dr. Michael Hast for donating the surgical drills used in this study. While no funding was accepted from Marrow Access Technologies for this study, Dr. Jason Koh, a co-author, serves as the Chief Medical Officer, and owns stock in this company.

## References

Bonadio MB, Friedman JM, Sennett ML, Mauck RL, Dodge GR, Madry H (2017) A retinaculum-sparing surgical approach preserves porcine stifle joint cartilage in an experimental animal model of cartilage repair. J. Exp. Orthop. 4: 0–4. doi:10.1186/s40634-017-0083-7.

Chen H, Sun J, Hoemann CD, Lascau-Coman V, Ouyang W, McKee MD, Shive MS, Buschmann MD (2009) Drilling and microfracture lead to different bone structure and necrosis during bone-marrow stimulation for cartilage repair. J. Orthop. Res. 27: 1432–1438. doi:10.1002/jor.20905.

Cole BJ, Farr J, Winalski CS, Hosea T, Richmond J, Mandelbaum B, De Deyne PG (2011) Outcomes after a single-stage procedure for cell-based cartilage repair: A prospective clinical safety trial with 2-year follow-up. Am. J. Sports Med. 39: 1170–1179. doi:10.1177/0363546511399382.

DiBartola AC, Everhart JS, Magnussen RA, Carey JL, Brophy RH, Schmitt LC, Flanigan DC (2016) Correlation between histological outcome and surgical cartilage repair technique in the knee: A meta-analysis. Knee 23: 344–349. doi:10.1016/j.knee.2016.01.017. http://dx.doi.org/10.1016/j.knee.2016.01.017.

Dyment NA, Jiang X, Chen L, Hong SH, Adams DJ, Ackert-Bicknell C, Shin DG, Rowe DW (2016) High-throughput, multi-image cryohistology of mineralized tissues. J. Vis. Exp. 2016: 1–11. doi:10.3791/54468.

Eldracher M, Orth P, Cucchiarini M, Pape D, Madry H (2014) Small subchondral drill holes improve marrow stimulation of articular cartilage defects. Am. J. Sports Med. 42: 2741–2750. doi:10.1177/0363546514547029.

Funk JF, Krummrey G, Perka C, Raschke MJ, Bail HJ (2009) Distraction osteogenesis enhances remodeling of remote bones of the skeleton: A pilot study. Clin. Orthop. Relat. Res. 467: 3199–3205. doi:10.1007/s11999-009-0902-y.

van Gaalen SM, Kruyt MC, Geuze RE, de Bruijn JD, Alblas J, Dhert WJA (2010) Use of fluorochrome labels in in vivo bone tissue engineering research. Tissue Eng. Part B. Rev. 16: 209–217. doi:10.1089/ten.teb.2009.0503.

Gao L, Goebel LKH, Orth P, Cucchiarini M, Madry H (2018) Subchondral drilling for articular cartilage repair: A systematic review of translational research. DMM Dis. Model. Mech. 11. doi:10.1242/dmm.034280.

Gobbi A, Nunag P, Malinowski K (2005) Treatment of full thickness chondral lesions of the knee with microfracture in a group of athletes. Knee Surgery, Sport. Traumatol. Arthrosc. 13: 213–221. doi:10.1007/s00167-004-0499-3.

Heijink A, Gomoll AH, Madry H, Drobnič M, Filardo G, Espregueira-Mendes J, van Dijk CN (2012) Biomechanical considerations in the pathogenesis of osteoarthritis of the knee. Knee Surgery, Sport. Traumatol. Arthrosc. 20: 423–435. doi:10.1007/s00167-011-1818-0.

Jafarzadeh SR, Felson DT (2018) Updated Estimates Suggest a Much Higher Prevalence of Arthritis in United States Adults Than Previous Ones. Arthritis Rheumatol. 70: 185–192. doi:10.1002/art.40355.

Kim IL, Pfeifer CG, Fisher MB, Saxena V, Meloni GR, Kwon MY, Kim M, Steinberg DR, Mauck RL, Burdick JA (2015) Fibrous Scaffolds with Varied Fiber Chemistry and Growth Factor Delivery Promote Repair in a Porcine Cartilage Defect Model. Tissue Eng. -Part A 21: 2680–2690. doi:10.1089/ten.tea.2015.0150.

Knutsen G, Drogset JO, Engebretsen L, Grøntvedt T, Isaksen V, Ludvigsen TC, Roberts S, Solheim E, Strand T, Johansen O (2007) A randomized trial comparing autologous chondrocyte implantation with microfracture: Findings at five years. J. Bone Jt. Surg. -Ser. A 89: 2105–2112. doi:10.2106/JBJS.G.00003.

Knutsen G, Isaksen V, Johansen O, Engebretsen L, Ludvigsen TC, Drogset JO, Grøntvedt T, Solheim E, Strand T, Roberts S (2004) Autologous Chondrocyte Implantation Compared with Microfracture in the Knee: A Randomized Trial. J. Bone Jt. Surg. -Ser. A 86: 455–464. doi:10.2106/00004623-200403000-00001.

Kraeutler MJ, Aliberti GM, Scillia AJ, McCarty EC, Mulcahey MK (2020) Microfracture Versus Drilling of Articular Cartilage Defects: A Systematic Review of the Basic Science Evidence. Orthop. J. Sport. Med. 8: 1–7. doi:10.1177/2325967120945313.

Kroell A, Marks P, Chahal J, Hurtig M, Dwyer T, Whelan D, Theodoropoulos J (2016) Microfracture for chondral defects: assessment of the variability of surgical technique in cadavers. Knee Surgery, Sport. Traumatol. Arthrosc. 24: 2374–2379. doi:10.1007/s00167-014-3481-8.

Madry H, Gao L, Rey-Rico A, Venkatesan JK, Müller-Brandt K, Cai X, Goebel L, Schmitt G, Speicher-Mentges S, Zurakowski D, Menger MD, Laschke MW, Cucchiarini M (2020) Thermosensitive Hydrogel Based on PEO–PPO–PEO Poloxamers for a Controlled In Situ Release of Recombinant Adeno-Associated Viral Vectors for Effective Gene Therapy of Cartilage Defects. Adv. Mater. 32: 1–8. doi:10.1002/adma.201906508.

Mainil-Varlet P, Van Damme B, Nesic D, Knutsen G, Kandel R, Roberts S (2010) A new histology scoring system for the assessment of the quality of human cartilage repair: ICRS II. Am. J. Sports Med. 38: 880–890. doi:10.1177/0363546509359068.

Martín AR, Patel JM, Zlotnick HM, Carey JL, Mauck RL (2019) Emerging therapies for cartilage regeneration in currently excluded ‘red knee’ populations. npj Regen. Med. 4. doi:10.1038/s41536-019-0074-7. http://dx.doi.org/10.1038/s41536-019-0074-7.

Meloni GR, Fisher MB, Stoeckl BD, Dodge GR, Mauck RL (2017) Biphasic Finite Element Modeling Reconciles Mechanical Properties of Tissue-Engineered Cartilage Constructs Across Testing Platforms. Tissue Eng. Part A 23: 663–674. doi:10.1089/ten.tea.2016.0191.

Miller RE, Grodzinsky AJ, Barrett MF, Hung HH, Frank EH, Werpy NM, McIlwraith CW, Frisbie DD (2014) Effects of the combination of microfracture and self-assembling peptide filling on the repair of a clinically relevant trochlear defect in an equine model. J. Bone Jt. Surg. - Am. Vol. 96: 1601–1609. doi:10.2106/JBJS.M.01408.

Mithoefer K, Mcadams T, Williams RJ, Kreuz PC, Mandelbaum BR (2009) Clinical efficacy of the microfracture technique for articular cartilage repair in the knee: An evidence-based systematic analysis. Am. J. Sports Med. 37: 2053–2063. doi:10.1177/0363546508328414.

Moore AC, DeLucca JF, Elliott DM, Burris DL (2016) Quantifying Cartilage Contact Modulus, Tension Modulus, and Permeability with Hertzian Biphasic Creep. J. Tribol. 138: 1–7. doi:10.1115/1.4032917.

Morisset S, Frisbie DD, Robbins PD, Nixon AJ, McIlwraith CW (2007) IL-1ra/IGF-1 gene therapy modulates repair of microfractured chondral defects. Clin. Orthop. Relat. Res.: 221–228. doi:10.1097/BLO.0b013e3180dca05f.

Murphy L, Helmick CG (2012) The Impact of osteoarthritis in the United States: A population-health perspective: A population-based review of the fourth most common cause of hospitalization in U.S. adults. Orthop. Nurs. 31: 85–91. doi:10.1097/NOR.0b013e31824fcd42.

Orth P, Duffner J, Zurakowski D, Cucchiarini M, Madry H (2016) Small-Diameter Awls Improve Articular Cartilage Repair after Microfracture Treatment in a Translational Animal Model. Am. J. Sports Med. 44: 209–219. doi:10.1177/0363546515610507.

Patel JM, Sennett ML, Martin AR, Saleh KS, Eby MR, Ashley BS, Miller LM, Dodge GR, Burdick JA, Carey JL, Mauck RL (2020) Resorbable Pins to Enhance Scaffold Retention in a Porcine Chondral Defect Model. Cartilage. doi:10.1177/1947603520962568.

Rahn BA, Perren SM (1971) Xylenol Orange, A Fluorochrome Useful in Polychrome Sequential Labeling of Calcifying Tissues. Stain Technol. 46: 125–129.

Riff AJ, Huddleston HP, Cole BJ, Yanke AB (2020) Autologous Chondrocyte Implantation and Osteochondral Allograft Transplantation Render Comparable Outcomes in the Setting of Failed Marrow Stimulation. Am. J. Sports Med. 48: 861–870. doi:10.1177/0363546520902434.

Ruehe B, Kershaw O, Niehues S, Nelson K (2008) Sudden death in miniature pigs. Lab Anim. (NY). 37: 65– 66. doi:10.1038/laban0208-65.

Schindelin J, Arganda-Carreras I, Frise E, Kaynig V, Longair M, Pietzsch T, Preibisch S, Rueden C, Saalfeld S, Schmid B, Tinevez J-Y, White DJ, Hartenstein V, Eliceiri K, Tomancak P, Cardona A (2012) Fiji: an open-source platform for biological-image analysis. Nat. Methods.

Sennett M, Friedman J, Ashley B, Stoeckl B, Patel J, Alini M, Cucchiarini M, Eglin D, Madry H, Mata A, Semino C, Stoddart3 M, Moutos FT, Estes BT, Guilak F, Mauck R, Dodge G (2021) Long Term Outcomes of Biomaterial-Mediated Repair of Focal Cartilage Defects in a Large Animal Model. Eur. Cells Mater. 41: 40–51. doi:10.22203/eCM.v041a04.

Sharma B, Fermanian S, Gibson M, Unterman S, Herzka DA, Cascio B, Coburn J, Hui AY, Marcus N, Gold GE, Elisseeff JH (2013) Human cartilage repair with a photoreactive adhesive-hydrogel composite. Sci. Transl. Med. 5: 1–10. doi:10.1126/scitranslmed.3004838.

Steadman JR, Briggs KK, Rodrigo JJ, Kocher MS, Gill TJ, Rodkey WG (2003) Outcomes of microfracture for traumatic chondral defects of the knee: Average 11-year follow-up. Arthrosc. - J. Arthrosc. Relat. Surg. 19: 477–484. doi:10.1053/jars.2003.50112.

Steadman JR, Rodkey WG, Briggs KK (2002) Microfracture to treat full-thickness chondral defects: surgical technique, rehabilitation, and outcomes. J. Knee Surg. 15: 170–176.

Theodoropoulos J, Dwyer T, Whelan D, Marks P, Hurtig M, Sharma P (2012) Microfracture for knee chondral defects: A survey of surgical practice among Canadian orthopedic surgeons. Knee Surgery, Sport. Traumatol. Arthrosc. 20: 2430–2437. doi:10.1007/s00167-012-1925-6.

Volz M, Schaumburger J, Frick H, Grifka J, Anders S (2017) A randomized controlled trial demonstrating sustained benefit of Autologous Matrix-Induced Chondrogenesis over microfracture at five years. Int. Orthop. 41: 797–804. doi:10.1007/s00264-016-3391-0.

Zanotto G, Liebesny P, Barrett M, Zlotnick H, Grodzinsky A, Frisbie D (2019) Trypsin Pre-Treatment Combined With Growth Factor Functionalized Self-Assembling Peptide Hydrogel Improves Cartilage Repair in Rabbit Model. J. Orthop. Res. 37: 2307–2315. doi:10.1002/jor.24414. http://dx.doi.org/10.1002/jor.24414.

Zedde P, Cudoni S, Giachetti G, Manunta ML, Masala G, Brunetti A, Manunta AF (2016) Subchondral bone remodeling: Comparing nanofracture with microfracture. An ovine in vivo study. Joints 4: 87–93. doi:10.11138/jts/2016.4.2.087.

